# Recently expanded clonal lineages of the rice blast fungus display distinct patterns of presence/absence of effector genes

**DOI:** 10.1101/2020.01.09.900308

**Authors:** Sergio M. Latorre, C. Sarai Reyes-Avila, Angus Malmgren, Joe Win, Sophien Kamoun, Hernán A. Burbano

## Abstract

**Background:** Understanding the mechanisms and timescales of plant pathogen outbreaks requires a detailed genome-scale analysis of their population history. The fungus *Magnaporthe* (Syn. *Pyricularia*) *oryzae* —the causal agent of blast disease of cereals— is among the most destructive plant pathogens to world agriculture and a major threat to the production of rice, wheat and other cereals. Although *M. oryzae* is a multihost pathogen that infects more than 50 species of cereals and grasses, all rice-infecting isolates belong to a single genetically defined lineage. Here, we combined multiple genomics datasets to reconstruct the genetic history of the rice-infecting lineage of *M. oryzae* based on 131 isolates from 21 countries.

**Results:** The global population of the rice blast fungus consists of a diverse set of individuals and three well-defined genetic groups. Multiple population genetic tests revealed that the rice-infecting lineage of the blast fungus probably originated from a recombining diverse group in South East Asia followed by three independent clonal expansions that took place over the last ∼200 years. Patterns of allele sharing identified a subpopulation from the recombining diverse group that introgressed with one of the clonal lineages before its global expansion. Remarkably, the four genetic lineages of the rice blast fungus vary in the number and patterns of presence/absence of candidate effector genes. In particular, clonal lineages carry a reduced repertoire of effector genes compared with the diverse group, and specific combinations of effector presence/absence define each of the pandemic clonal lineages.

**Conclusions:** Our analyses reconstruct the genetic history of the rice-infecting lineage of *M. oryzae* revealing three clonal lineages associated with rice blast pandemics. Each of these lineages displays a specific pattern of presence/absence of effector genes that may have shaped their adaptation to the rice host and their evolutionary history.

## Background

Plant diseases are a persistent threat to food production due to a notable increase in the emergence and spread of new pathogens [1, 2]. Understanding the mechanisms and timescales associated with new epidemics is essential for both basic studies and the implementation of effective response measures [3]. A fundamental component of this knowledge is a detailed genome-scale understanding of the population structure and dynamics of global plant pathogen populations [4–6]. Population genetic information drives the selection of isolates for activities as diverse as basic mechanistic research and plant germplasm screening for disease resistance. It also helps to pinpoint the origin of pandemic strains and the evolutionary potential of different pathogen populations [7–12]. A thorough understanding of the global population structure is essential for any surveillance program that aims at rapidly detecting pathogen incursions into new geographical areas. In addition, the recent knowledge gained in the biology of pathogen effectors —secreted molecules that modulate host responses— brings yet another dimension to the population genetics framework, as it enables the reconstruction of the evolutionary history of virulence traits and helps guide the deployment of disease resistant cultivars [7, 13–16].

Fungal plant pathogens account for ∼10-80% of crop losses in agriculture and are viewed as a major threat to global food security [1, 2, 17, 18]. Cereal crops like rice, oat, millet, barley and wheat have provided the foundation of modern agriculture and the success of humankind. Today’s agriculture is facing the challenge of ensuring global food security for an ever-expanding world population, which is estimated to exceed 9 billion within the next 30 years [19]. The fungus *Magnaporthe* (Syn. *Pyricularia*) *oryzae*, the causal agent of blast disease of cereals, is often ranked as the most destructive fungal pathogen, causing losses in rice production that, if mitigated, could feed several hundred million people [1, 20]. Despite its Linnean name, *M. oryzae* is a multihost pathogen that can also cause the blast disease on other cereal crops, notably on wheat where it has recently spread from South America to Bangladesh resulting in destructive outbreaks [8, 21, 22].

Comparative genomics analyses provided insights into the population structure and host-specialization of *M. oryzae* [23–25]. This pathogen consists of a complex assemblage of genetically distinct lineages that tend to be associated with particular host genera [25]. Remarkably, all rice-infecting isolates belong to a single genetic lineage that is thought to have originated from isolates infecting foxtail millet *(Setaria italica* and *Setaria viridis*). *M. oryzae* host-specific lineages exhibit limited gene flow but recurrent gene gain/loss particularly in regions of the genome linked to transposable elements [23, 24]. As in many other plant pathogens, effector genes exhibit a high degree of presence/absence polymorphisms and signatures of adaptive evolution (e.g. higher rate of non-synonymous over synonymous mutations) [24]. Loss of so-called AVR effector genes —activators of host immunoresponses— can dramatically impact the fitness of the blast fungus by enabling virulence on resistant host genotypes [22, 26, 27].

Although the genome sequence of the *M. oryzae* strain 70-15 was at the time of its publication the first fungal plant pathogen genome to be described [28], it took about a decade before comparative genomics analyses of this pathogen started to be reported [23, 24, 29]. Until recently, understanding of the population genomics structure of the rice blast fungus has remained limited. In 2018, two studies reported whole genome sequences from non-overlapping sets of globally distributed rice-infecting *M. oryzae* isolates [30, 31]. Both studies suggested the presence of a diverse Southeast Asian population and two major clonal groups. However, due to sampling or analytical limitations the two studies reached different conclusions about the composition of worldwide populations.

Here, we performed a combined analysis that builds on the studies of Gladieux *et al*. [30] and Zhong *et al*. [31] to reconcile the two datasets and increase the number of examined *M. oryzae* individuals to 131 isolates from 21 countries. This has enabled us to assess the global genetic structure of rice-infecting *M. oryzae* more comprehensively than the prior separate analyses of the two datasets. We discovered that the global population of the rice blast fungus consists of a diverse set of individuals and three well-defined genetic groups. Multiple population genetic tests revealed that the rice blast fungus probably originated from a recombining population in South East Asia followed by three independent clonal expansions that took place over the last ∼100-200 years. Patterns of allele sharing identified a subpopulation from the recombining group that introgressed with one of the clonal lineages before its global expansion. Remarkably, the genetic lineages of the rice blast fungus vary in the number and patterns of presence/absence of secreted protein predicted as effectors. In particular, the clonal lineages are defined by specific sets of effectors that may have shaped their adaptation to the rice host and their evolutionary history.

## Results and discussion

### The global population structure of rice-infecting *Magnaporthe oryzae* consists of three well defined genetic groups and a diverse set of individuals

To assess the global population structure of rice-infecting *M. oryzae*, we used a total of 131 genome sequences from Gladieux *et al*. (N=43) [30] and Zhong *et al*. (N=88) [31]. The combined use of samples from these two studies increases not only the number of *M. oryzae* samples but also their geographical spread (Supp. Table 1 and Fig. 2B-C). We identified a total of 39,862 Single Nucleotide Polymorphism (SNPs) (see the “Methods” section). For subsequent analyses, we only used SNPs ascertained in all samples (“full information”) (N=11,478 SNPs).

**Figure 1.**
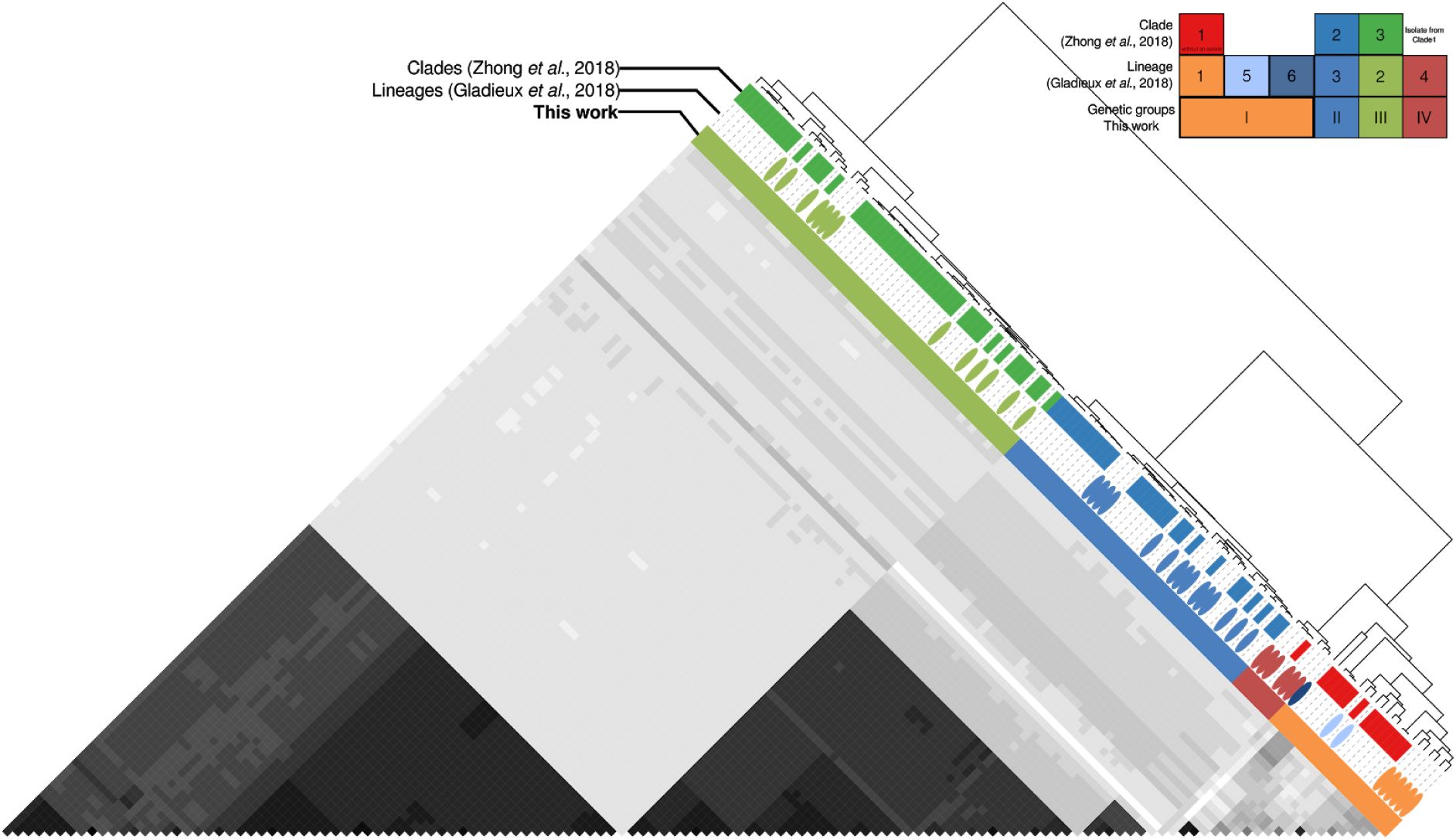
Genetic clustering of *Magnaporthe oryzae* reveals three defined groups and a diverse set of individuals. The pairwise relatedness between *M. oryzae* samples (X and Y) was estimated using f3-outgroup statistics of the form *f3*(X, Y; Outgroup), which measures the amount of shared genetic history (genetic drift) between X and Y after the divergence from an outgroup (*M. oryzae* strain from wheat). The hierarchical clustering is based on *f3-scores* resulting from f3-outgroup statistic calculations. Darker colors indicate more shared drift.

**Figure 2.**
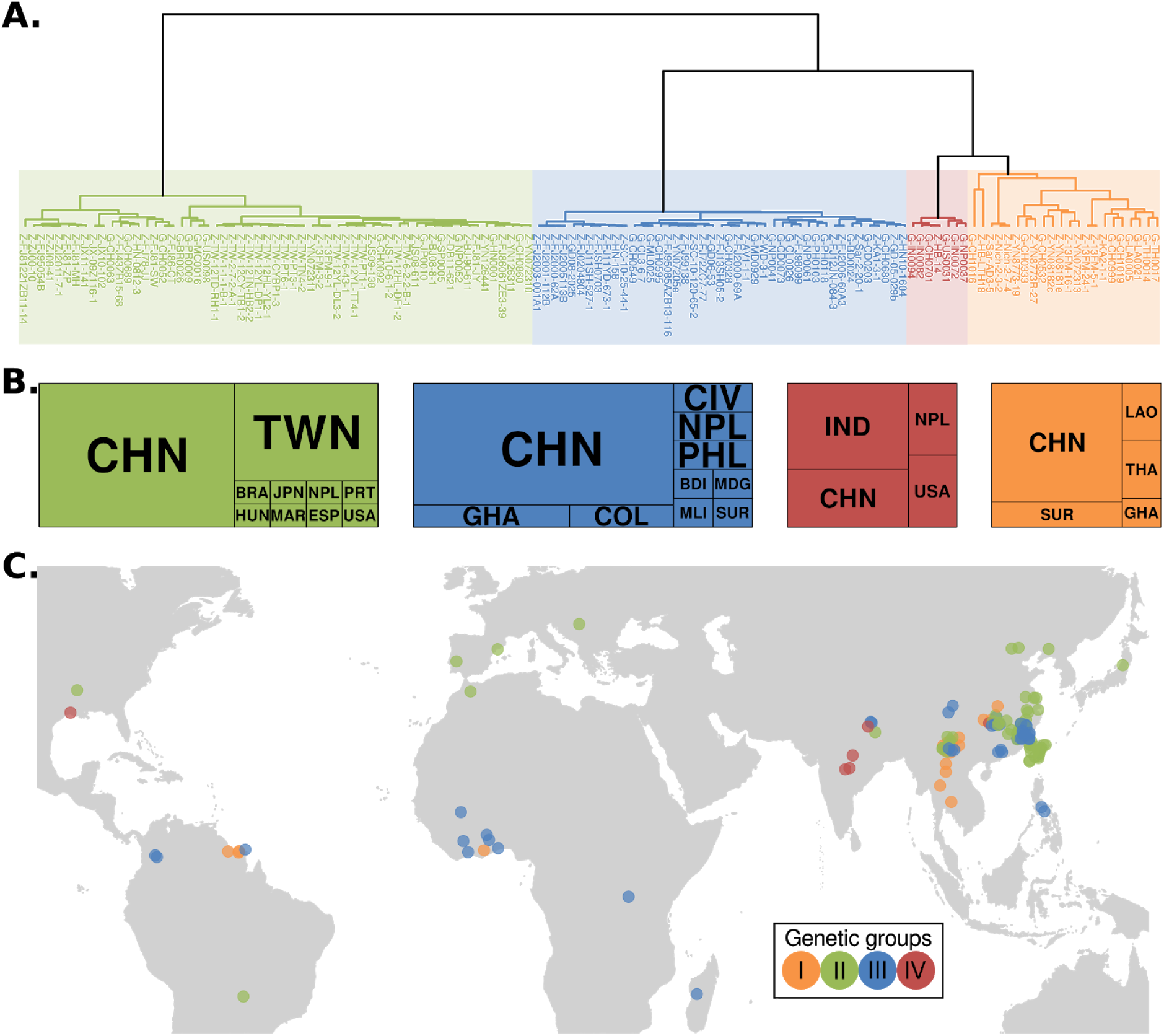
Geographic location of *Magnaporthe oryzae* isolates shows global distribution of defined genetic groups (II-III) and a preferential South-East Asian location for the diverse group (I). **(A)** Dendrogram showing the hierarchical clustering based on pairwise f3 values (same as Fig. 1). Prefixes of the isolates names correspond to the database source: G = Gladieux *et al*., 2018 [30]; Z = Zhong *et al*., 2018 [31]. (**B**) Country of origin for *M. oryzae* isolates. The overall size of the boxes represents the total number of samples within each genetic group. The size of each internal box is proportional to the number of isolates per country. Countries are represented as three-letter codes (ISO 3166-1 alpha-3): BDI=Burundi, BRA=Brazil, CHN=China, CIV=Côte d’Ivoire, COL=Colombia, ESP=Spain, GHA=Ghana, HUN=Hungary, IND=India, JPN=Japan, LAO=Lao People’s Democratic Republic, MAR=Morocco, MDG=Madagascar, MLI=Mali, NPL=Nepal, PHL=Philippines, PRT=Portugal, SUR=Suriname, THA=Thailand, TWN=Taiwan, Province of China, USA=United States of America. **(C)** Geographical origin of samples used in this study. A random jitter was used on the coordinates of geographical-close samples for better visualization.

We first sought to investigate the number of distinct genetic groups in our global sample of *M. oryzae* given previous discrepancies in the number of clades or lineages identified in the two studies. We identified three well-defined groups and a diverse set of individuals based on two lines of evidence. First, we used f3-outgroup statistics [32] to evaluate the pairwise relatedness between *M. oryzae* samples relative to an outgroup. The f3-outgroup statistics measure the amount of shared evolutionary history between samples, which can be interpreted as shared genetic drift (always relative to an outgroup). We summarized the results of all tests by performing hierarchical clustering based on pairwise shared genetic drift comparisons, i.e. z-scores derived from f3-outgroup statistic tests (Fig. 1). Additionally, we calculated pairwise Hamming genetic distances between all samples and summarized the information using Principal Component Analysis (PCA). The samples clustered again in three distinct groups and one diverse set of individuals using PC1, 2 and 3, which together explained more than 90% of the variance (Supp. Fig. 1A). We assessed the robustness of these clusters using Silhouette scores, which indicate how similar an individual is to its own cluster compared to other clusters [33]. We found that the best mean Silhouette scores were obtained when the dataset was divided into four clusters (Supp. Fig. 1B).

Since our two approaches consistently revealed the presence of four groups, we named them group I, II, III and IV. Whereas group II and III are geographically widespread, Group I is mainly located in South-East Asia and group IV in the Indian subcontinent (Fig. 2). The correspondence between our classification and previously described nomenclatures can be found in Supp. Table 2. Our grouping recapitulates the four lineages proposed by Saleh *et al*. based on microsatellite data [34]. Zhong *et al*. [31] divided their dataset in three groups (I-III) but did not identify group IV, since their dataset only include one individual from this group. In addition to groups I-IV, Gladieux *et al*. [30] identified two additional lineages based on a set of phylogenetic analyses. The combined analysis presented here showed that these additional lineages from Gladieux *et al*. are within the genetic diversity of group I, thus splitting of group I is not warranted.

### Global population of rice-infecting *Magnaporthe oryzae* probably arose from a recombining South East Asian population followed by clonal expansions

To determine the evolutionary origin of the four *M. oryzae* groups identified in this study, we used a set of statistics that evaluate genetic diversity, recombination and population differentiation. Initially, we visualized the relationships among samples using a phylogenetic network, which are more appropriate for visualizing reticulate evolution (Fig. 3A) [35]. We found that group I exhibited a high degree of reticulation. In contrast, the phylogenetic network showed long internal branches with terminal star-shape phylogenetic configurations almost devoid of reticulations for the well-defined groups II, III and IV (Fig. 3A). Such configurations are typical of expanding populations after genetic bottlenecks, driven, for instance, by clonal expansions [36]. We, therefore, queried whether genetic diversity levels and recombination rates support clonality in groups II, III, and IV. Two lines of evidence support clonality in these groups compared with the diverse group I: i) reduced nucleotide diversity measured as pi (*π*) [37] (Fig. 3B); ii) lower detectable recombination events calculated using the four-gamete test [38] (Fig. 3C). The reduced levels of diversity in groups II, III, and IV in conjunction with their star-like phylogenies are tell-tale signs of populations that have experienced a strong reduction of diversity followed by a population expansion. To calculate a proxy for recombination we used the four-gamete test, which puts a bound to the minimum number of recombination events in a sample [38]. Although it is known that this test underestimates recombination events, it is a simple and useful proxy for differences in recombination between populations. Our results showed that groups II, III and IV have on average ∼5-fold less recombination events than the diverse group I. In agreement with our analysis, metainformation obtained by Gladieux *et al*. and Zhong *et al*. [30, 31] showed that in almost all cases only one mating type was present in groups II, III and IV, whereas the two mating types were segregating in the diverse group I (Supp. Table 1 and Supp. Fig. 3). In sum, we conclude that group II, III and IV are likely clonal lineages, while group I consists of genetically diverse and recombining individuals (Fig. 3A-C). The original microsatellite-based study by Saleh *et al*. [34] reported a high level of genetic variability in group IV, however both our analyses and the ones carried out by Gladieux *et al*. [30] supported the clonal nature of this group.

**Figure 3.**
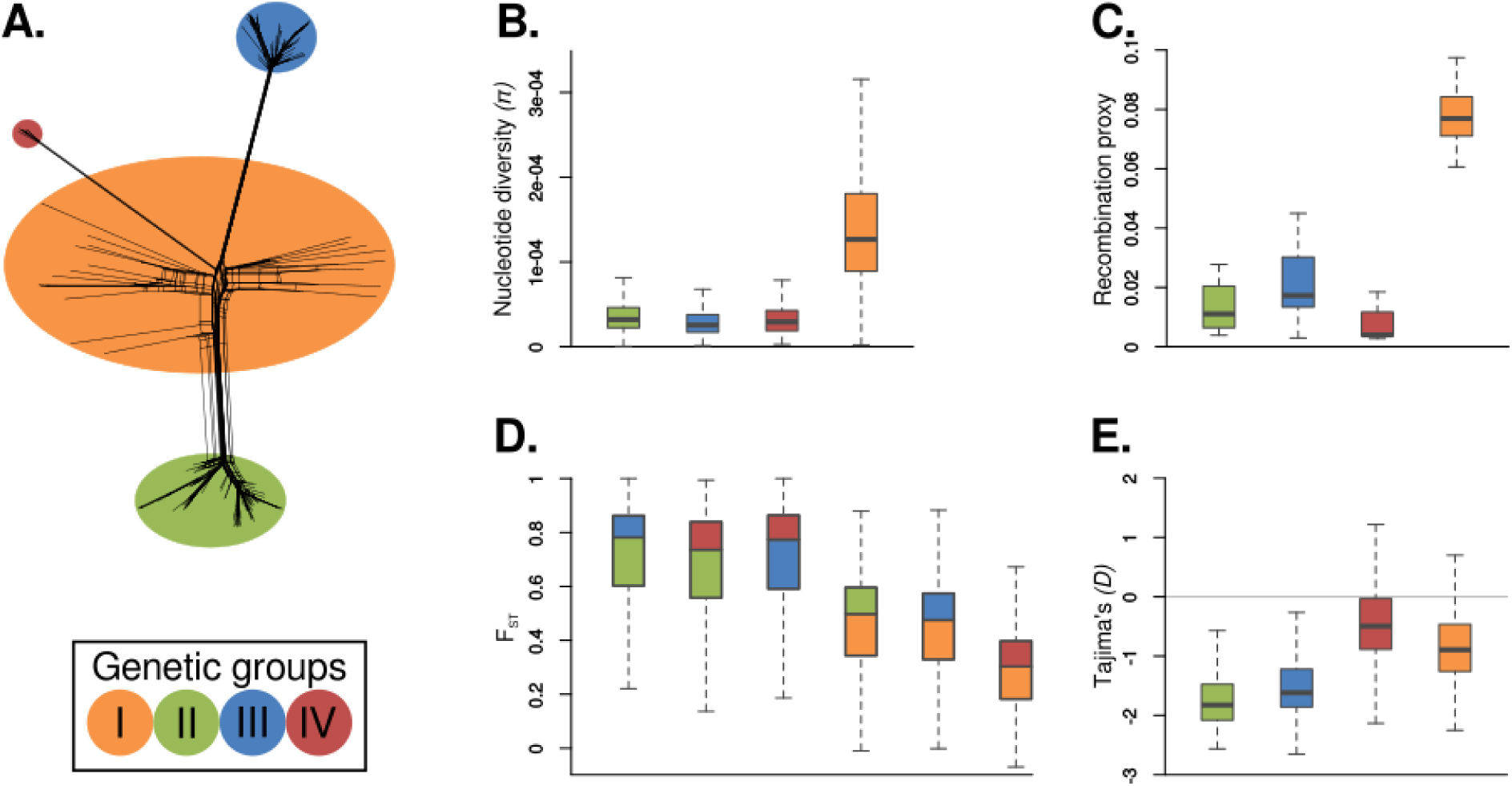
*Magnaporthe oryzae* population structure is driven by local recombination and global clonal expansions. **(A)** Phylogenetic network showing the three well-defined groups (green, blue and red) and the diverse set of individuals (orange) from figure 1. **(B)** Within-population comparisons of nucleotide diversity measured as π. **(C)** Recombination proxy calculated by dividing the number of violations of the Four-gamete Test by the total number of SNPs. **(D)** Genetic distances between groups measured as Fixation Indices (F_st_). The box colors depict the pairwise comparisons between groups. **(E)** Tajima’s *D*.

To further investigate the relationships and demographic history of *M. oryzae* groups, we measured population differentiation among groups and leveraged the allele frequency spectrum for each group individually. To measure population differentiation we used F_ST [39]_ and found that when clonal groups II, III, and IV are compared among them, their F_ST_ distances were the highest, reflecting a long history of independent drift. In contrast, whenever the diverse group I is compared with any of the clonal groups, the F_ST_ distances decreased, suggesting that group I is a common source of genetic diversity for all clonal lineages (Fig. 3D). Subsequently, for every group we investigated their corresponding allele frequency spectrum using Tajima’s *D [40]*, as this statistic records changes in allele frequencies driven, for instance, by variation in population sizes. We found that Tajima’s *D* values for all clonal lineages were negative (Fig. 3E). A demographic interpretation of negative Tajima’s *D* values is consistent with population bottlenecks followed by population expansions and a concurrent accumulation of rare alleles. Negative Tajima’s *D* values are consistent with star-like phylogenies, as new mutations that occurred post-bottleneck accumulate in terminal branches lowering Tajima’s *D* values. One possibility is that the difference in average Tajima’s *D* values between the clonal groups is due to different timings of clonal expansions, *i*.*e*. group IV clonal expansion might have taken place later than group II and III. Overall, our results are consistent with a model where south-east Asia is a likely centre of origin of rice-infecting *M. oryzae* and in which three distinct clonal lineages arose from this ancestral population. These findings are consistent with previous models of the evolution of the rice lineage of *M. oryzae* [34].

### *Magnaporthe oryzae* rice-infecting clonal lineages are estimated to have arisen in the last 200 years

To estimate the divergence time of the clonal expansions of *M. oryzae*, we used Bayesian phylogenetic analysis leveraging the sample collection dates for tip-calibration [41, 42]. To carry out the analysis, we first removed the diverse group I and used only the three clonal lineages, as the recombining group violates the assumptions of phylogenetic reconstruction. We estimated an evolutionary rate of 1.32 e-8 substitutions/site/year (1.17 e-8 - 1.47 e-8 HPD 95%), which was similar but lower than a previously calculated rate (1.98e-8 substitutions/site/year) [30]. Our approach of including only the clonal lineages permitted the reconstruction of a robust phylogeny and a more accurate estimation of divergence times, as reflected in the high posterior probabilities supporting the nodes and the narrow HPD 95% confidence intervals of node ages (Fig. 4). This contrasts with previous studies that included individuals from the diverse recombining group I in the phylogenetic analysis and produced broader HPD 95% confidence intervals (Gladieux et al., 2018; Supp. Fig. 3 [30]).

**Figure 4.**
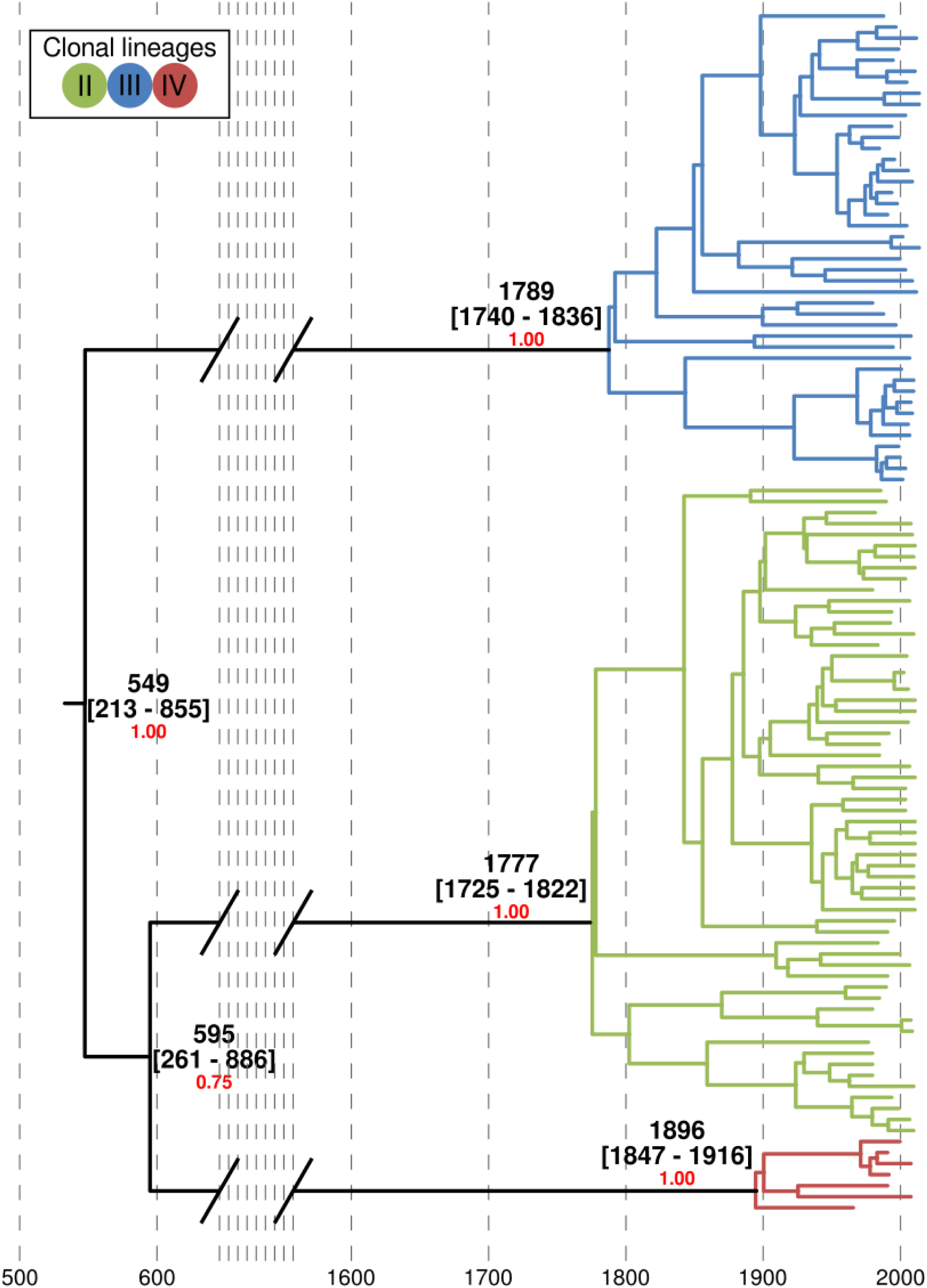
Clonal expansions of *Magnaporthe oryzae* took place in the last 200 years. Bayesian tip calibrated phylogenetic tree using individuals belonging to clonal lineages. Average, and HPD 95% confidence intervals are shown in calendar years. The Bayesian posterior probability is shown in red for nodes leading to the clonal lineage expansions.

The phylogenetic reconstructions revealed that all three clonal expansions occurred relatively recently over the last 200 years (123-242) (Fig. 4). In the previous section, we hypothesized that differences in average Tajima’s *D* values between the clonal groups is due to different timings of clonal expansions, i.e. group IV might have had less time to accumulate mutations in the terminal branches, which resulted in less negative Tajima’s *D*. The phylogenetic analyses are consistent with this hypothesis showing that the less negative Tajima’s *D* of group IV —compared to group II and II— corresponds to a more recent clonal expansion of Group IV.

### Patterns of allele frequency sharing identify introgression between a subpopulation of the diverse group I and clonal lineage II

Since the identification of admixture between populations facilitates the reconstruction of the evolutionary history of populations, we investigated the admixture history of *M. oryzae* using *D-statistics* [43, 44]. This test employs counts of site patterns, which are patterns of alternative alleles at a given genomic position and evaluates whether these site patterns support one of two alternative discordant topologies. The *D*-statistics will return a value of zero if the two discordant phylogenies are supported equally, whereas positive or negative values indicate asymmetric support and, therefore, introgression. We test the three possible configurations of the following form: *D*(Outgroup, Diverse Group I; Clonal lineage X, Clonal lineage Y) (tree insets in Fig. 5A-C). Whilst for clonal lineages II, III, and VI we used a strain representative for each clonal lineage, we performed a test for every one of the 22 members of the diverse group I. The test will retrieve positive values when the diverse group I is closer to clonal lineage Y and negative values when the diverse group I is closer to clonal lineage X. We found that group II has drifted farther apart from the diverse group I than the two other clonal lineages, as manifested from positive *D*-statistics when group II was included (as clonal lineage X) in the comparisons (Fig. 5B-C). This accumulation of genetic drift is consistent with the fact that group II was the clonal lineage that diverged earliest from the recombining diverse group (Fig. 4). We retrieved positive *D-*statistics in tests including almost all individuals of the diverse group I, with the exception of two individuals collected in China —*CH1016* [30] *and HB-LTH18* [31]*— that showed strong signals of genetic introgression with the clonal lineage II, as manifested by negative D*-statistic values (Fig. 5B-C). Since we detected introgression between these two Chinese samples and all members of group II regardless of their geographic origin (Supp. Fig. 4A-B), we inferred that the admixture should have taken place before the clonal expansion that gave rise to group II about 197-294 years ago (Fig. 4). Previous attempts to detect interlineage recombination were not statistically robust and plagued with false positives [30]. In contrast, *D*-statistics provide a statistically robust framework that reliably permits distinguishing between introgression and incomplete lineage sorting using genome-wide SNPs [43, 44].

**Figure 5.**
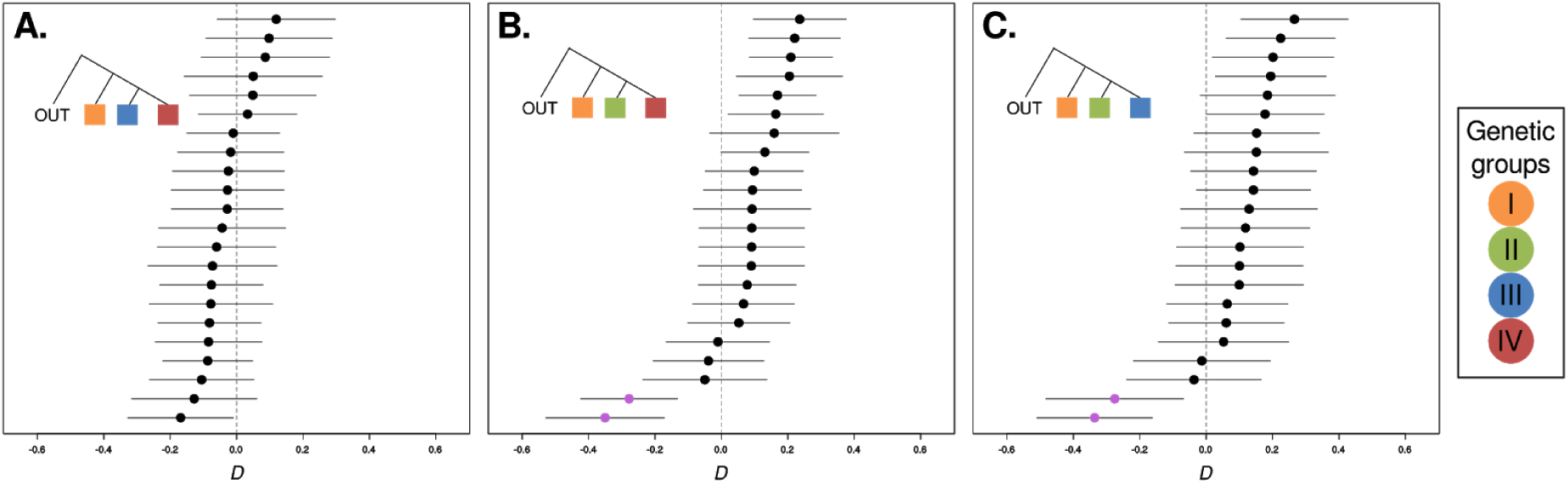
Patterns of allele frequency sharing identify introgression between a Chinese *Magnaporthe oryzae* subpopulation and clonal lineage II. *D*-statistics using three different phylogenetic configurations (depicted as colored inset trees). **(A)** *D*(Outgroup, Orange; Blue, Red). **(B)** *D*(Outgroup, Orange; Green, Red). (**C**) *D*(Outgroup, Orange; Green, Blue). In all cases, a *M. oryzae* strain from wheat was used as an outgroup and a fixed individual was selected as representative from each clonal lineage (Blue, Orange, Red). Points represent *D*-statistic tests for each of the 22 individuals assigned to the diverse clade (Orange), and lines depict 95% confidence intervals. Purple dots in **B**. and **C**. correspond to Chinese individuals *CH1016* and *HB-LTH18*, which are the closest individuals to the Green clonal lineage.

To further investigate the extent and location of the introgression between group II and the two Chinese Group I individuals (*CH1016/HB-LTH18*), we segmented the genomes of the two Chinese individuals based on their similarity at segregating sites to either Group I or Group II (Supp. Fig 5B). This analysis revealed that the genome-wide percentage of Group II-like fragments in the Chinese individuals is 44.58%, including a ∼4 Mb region in chromosome 3 (Supp. Fig. 5B). To test whether those fragments are a good proxy for the percentage of introgression we carried out two additional tests. First, we repeated the *D*-statistic test presented in figure 5B and supplementary figure 5A, but this time removing the candidate introgressed fragments. In contrast to the outcome of the test with whole-genome data, this time the test did not indicate introgression, i.e., it was not different from zero (Supp. Fig 5C). Second, we estimated the proportion of introgression by using a f4-ratio test [45] with the following setup: (Group III, Group II, Group I (without introgressed Chinese individuals), Outgroup) / (Group III, Group II, Chinese introgressed individuals, Outgroup). This test estimated the mixture proportion to be ∼31.68%, a lower but similar value to the overall percentage of identified Group II-like fragments in the Chinese individuals.

### The emergence of rice-infecting *Magnaporthe oryzae* coincides with rice domestication

Estimating the date of emergence of crop pests is fundamental for understanding how different factors, such as the agricultural practices associated with the neolithic revolution, have shaped the disease landscape of modern crops. Previous analyses [30, 31] have attempted to date the emergence of rice-infecting *M. oryzae* using phylogenetic approaches that rely on the collection dates of the isolates to calibrate the inferred phylogeny [46]. Given that those previous studies [30, 31] included the recombining Group I, which is not suitable for phylogenetic analysis, and produced broad confidence intervals for divergence times [30], we revisited the dating of the emergence of rice-infecting *M. oryzae*. Since *Setaria*-infecting *M. oryzae* strains are the closest genetic group to rice-infecting individuals [47], we used a *Setaria*-infecting isolate together with the samples from the different clonal lineages to calculate the divergence time of rice-infecting *M. oryzae*. We followed the same Bayesian phylogenetic analysis approach described before for the dating of the emergence of the *M. oryzae* clonal lineages. Our analysis suggested a host shift from millet to rice dated 9,503 Years Before Present (YBP) (11,867 - 7,407 YBP HDP 95%) (Fig. 6). This estimation overlaps with the archaeobotanical evidence that suggests a domestication period between 12,000 to 7,000 YBP [48] and the spread of domesticated varieties between 6,900 to 6,600 YBP [49]. Indeed, the coincidence between rice domestication and emergence of rice-infecting *M. oryzae* has been suggested already with and without strong confidence on the reported intervals, 8,000 to 12,000 YBF [31] and 1,200 to 21,700 YBF [30], respectively. Our analysis supports the hypothesis of a simultaneous process between shifts in human crop domestication and pathogen adaptation via host shift.

**Figure 6.**
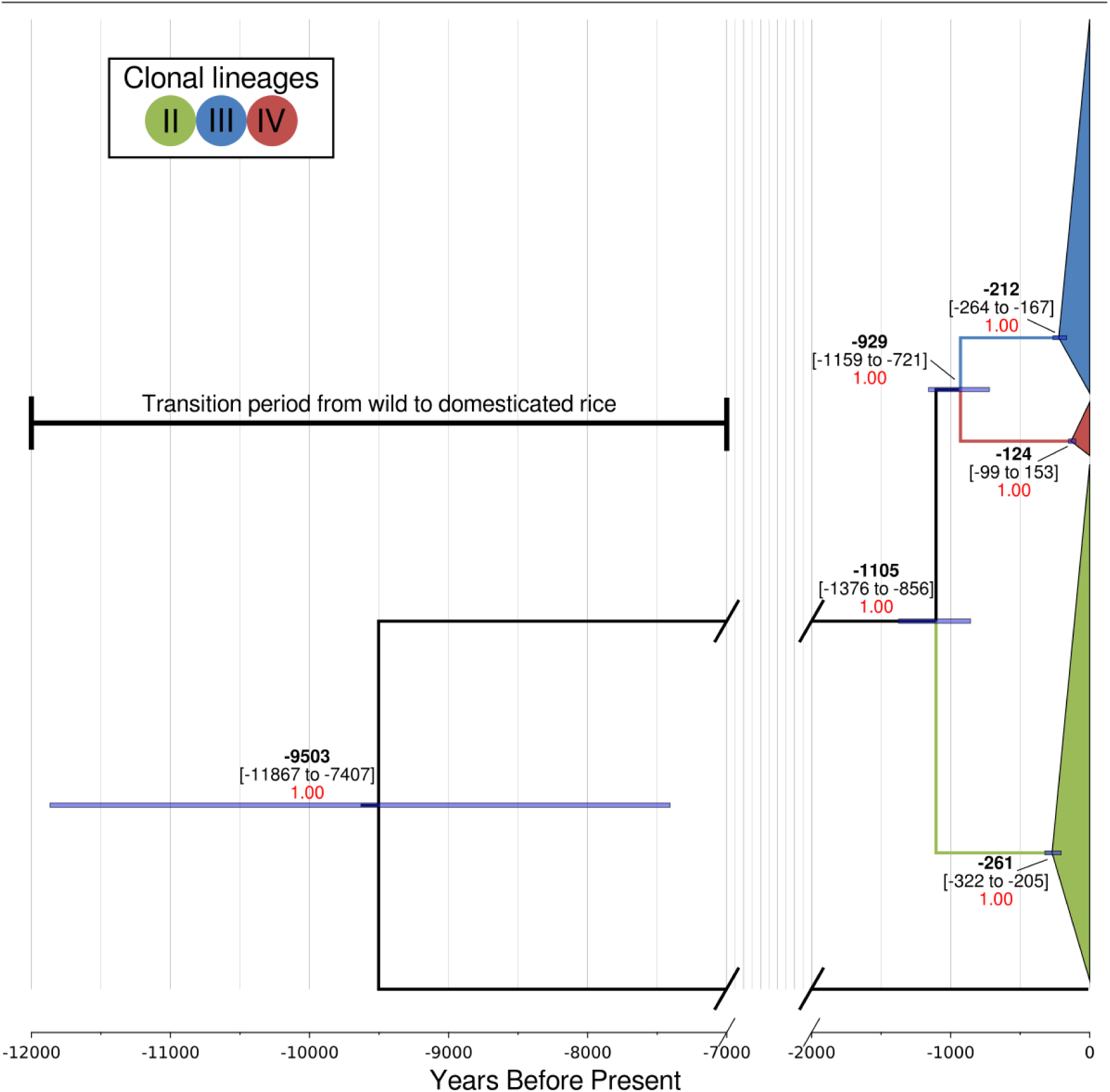
The emergence of rice infecting *Magnaporthe oryzae* overlaps with rice domestication. Bayesian tip calibrated phylogenetic tree using individuals belonging to clonal lineages and a *Setaria*-infecting individual as outgroup. Average time estimates and their HPD 95% confidence intervals are shown in years. Blue horizontal bars represent HPD 95% at the nodes. The Bayesian posterior probability is shown as numbers in red.

### Lineages of *Magnaporthe oryzae* show distinct patterns of presence/absence of effector genes

In *M. oryzae*, effector genes exhibit a high rate of structural variation as illustrated by presence/absence polymorphisms [24]. We investigated the distribution of known and predicted effector genes within the population structure framework we defined for the rice lineage of *M. oryzae*. We mapped the genome sequences of the 131 isolates to the sequences of 178 known and candidate effectors predicted from the genomes of *M. oryzae* from hosts as diverse as rice, wheat, finger millet, foxtail millet, oat and *Digitaria* spp. [50]. This pan-effectorome set enabled us to capture as much effector gene diversity as possible. In total, 134 effectors were identified in the 131 isolates (Supp. Table 3). Remarkably, the number of effectors per isolate varied from 108 to 125 with clonal lineages carrying a reduced repertoire of effector genes compared with the diverse genetic group (Fig. 7A-B). This indicates that clonal-expansion-driven bottlenecks not only reduced the overall genetic diversity of all pandemic clonal lineages but also their repertoire of dispensable genes such as effectors. In pathogenic bacteria, a reduction in the effectiveness of purifying selection has been associated with an increase in gene loss [51]. Moreover, gene loss is particularly prevalent in clonal pathogenic bacteria and has been postulated as a source of phenotypic variation in these otherwise genetically similar species [52]. The association between gene loss and reduced purifying selection in bacteria is a consequence of their strong deletional bias, i.e. bacteria with reduced effective population size experience genome reduction [53]. In contrast, eukaryotes with small effective population sizes have larger genomes [54]. This relation is, however, more complex in the rice blast fungal phylum Ascomycota, where both genome expansions and reductions have been observed [55]. It remains to be tested whether the concurrent loss of genetic diversity and dispensable/non-core genes is a widespread consequence of clonality-driven bottlenecks in eukaryotic pathogens that can reproduce both sexually and asexually.

**Figure 7.**
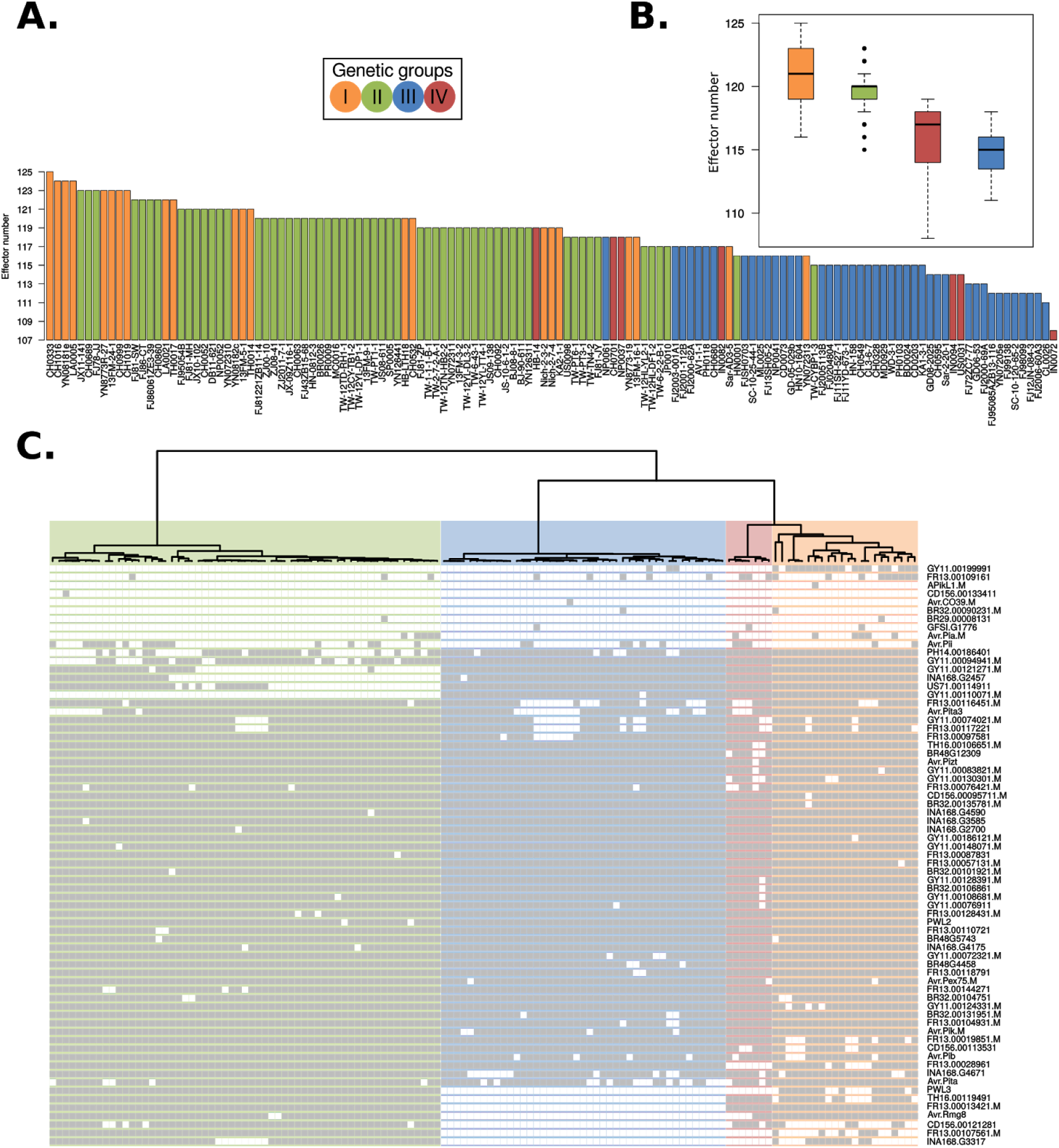
Rice blast genetic lineages vary in the number and patterns of presence/absence of candidate effector genes. **(A)** Clonal lineages carry a reduced repertoire of effector genes compared with the diverse group I. The box-and-whisker plots show the distribution of effector number per isolate for each genetic group. (**B**) The dendrogram shows the clustering based on f3-outgroup statistic (as in Fig. 1). Light and dark colors on the rows, represent absence and presence of effectors, respectively. Rows were grouped using a hierarchical clustering algorithm.

We next mapped the distribution of the subset of 69 effectors that display presence/absence polymorphisms across all strains (Fig. 7C). The resulting matrix clearly shows that there are distinct patterns of presence/absence of effectors across the genetically defined groups. For example, a set of four effectors (Avr-Rmg8, FR13.00013421, TH16.00119491 and CD156.00121281) are absent in Group III. Likewise, PWL3, INA168.g3317 and FR13.00107561.M are absent in Groups III and IV, GY11.00110071.M is absent in Group II, and FR13.00028961 is absent in Group IV (Fig. 7C, Supp. Table 4).

To determine which effectors have the strongest association with the defined genetic structure, we conducted two separate analyses based on the presence/absence effector repertoire per isolate. First, a PCA and effector loadings analysis revealed a set of 13 effectors that explained 90% of the variance of both PC1 and PC2 (Supp. Fig. 6A-B). Similarly, by using extremely randomized trees (a classification machine learning technique), we identified a set of 16 effectors that explained 90% of the variance (Supp. Fig. 6C). Although the two methods produced different rankings of the impact of each effector genes, we found an overlap of 92.3% between the top 13 effectors found in the two subsets. In both cases the top effectors reproduced the separation of the isolates in the described genetic clusters (Supp. Fig. 6D-E). A close inspection of this group of top effectors, which were selected in an unbiased way, revealed that they are differentially (almost) present or (almost) absent in the four *M. oryzae* genetic groups (Supp. Table 4). Thus, this group of effectors might have played an important role in the initial adaptation of *M. oryzae* clonal expansions to different rice subspecies and varieties.

The matrix in Fig. 7C indicates that patterns of presence/absence of effector genes reflect different timescales in the evolution of the clonal lineages of *M. oryzae*. AVR effectors, such as AVR-Pia and AVR-Pii, show a patchy distribution within the clonal lineages consistent with the view that they have been recurrently deleted in *M. oryzae* populations to generate virulent races [26]. This may reflect the fact that their matching resistance genes have been repeatedly bred and deployed into rice cultivars. Other candidate effectors that display a similar patchy distribution may be candidate AVR effectors that are detected by one of the dozens of blast resistance genes that have been bred into rice cultivars.

Our finding that the clonal lineages of rice-infecting *M. oryzae* display distinct repertoires of effectors raises a number of interesting questions. It is possible that this reflects the distinct genotype of the founding individual of the given clone. It is also possible that the absence of a given AVR effector(s) has facilitated the spread of the clonal lineage to otherwise resistant host genotypes as previously noted in *M. oryzae [22, 29, 56, 57]*. In the future, it would be interesting to test the extent to which effectors that define the clonal lineages are detected by particular resistance genes. For example, AVR-Rmg8, which is known in wheat blast isolates to mediate avirulence on Rmg8 containing wheat varieties, may also be detected by a rice resistance gene. Future experiments will tease out the degree to which the distinct effector repertoires of the clonal lineages of *M. oryzae* reflect their adaptation to the rice host and their evolutionary history. Such analyses will require new genomic resources that permit a more accurate identification of effectors in canonical chromosomes and mini-chromosomes [58]. To this aim, it will be fundamental to generate multiple reference genomes sequenced with long-read technologies in conjunction with a detailed characterization of structural variation and genomic rearrangements, which will include a per isolate inventory of mini-chromosome repertoires.

## Conclusion

Our analyses reconstruct the genetic history of the rice-infecting lineage of *M. oryzae* revealing three clonal lineages that have emerged over the last ∼100-200 years and have been associated with rice blast pandemics. These lineages display specific patterns of presence/absence of effector genes that may have shaped their adaptation to the rice host and their evolutionary history. These findings provide a framework for further comparative analyses of the genomes of rice-infecting *M. oryzae*. One particular interesting research avenue will be to establish the degree to which structural variation, notably mini-chromosomes, have impacted the evolution of this lineage.

## Methods

### Datasets and mapping

We used *M. oryzae* Illumina reads from two recent resequencing studies (43 samples from Gladieux *et al*. [30], and 88 samples from Zhong *et al*. [31] (Supp. Table 1)). Raw sequencing reads were downloaded and mapped to the *M. oryzae* reference genome (*GUY-11* PacBio assembly [59]) using *bwa-mem* V.0.7.12 [60] with default parameters.

### Variant identification and filtering

De novo variants were identified using *GATK* V.3.8.0 [61]. The following set of filters were applied: QD < 5.0; QUAL < 5000.0; MQ < 20.0; −2.0 < ReadPosRankSum < 2.0; −2.0 < MQRankSum < 2.0; −2.0 < BaseQRankSum < 2.0. In all subsequent analysis we used only biallelic SNPs present in all samples (“full information”).

### Population Structure Analyses

To assess the global population structure of *M. oryzae* we first determined patterns of allele sharing using f3-outgroup statistics [32]. We performed the test using the program *qp3Pop* from the *AdmixTools* package [45]. The test was used to establish the pairwise relatedness between *M. oryzae* samples (X and Y) after divergence from an outgroup: *f3*(X, Y; Outgroup). We used a deeply diverged *Setaria*-infecting *M. oryzae* strain SA05-144 [24] as outgroup. We calculated z-scores for every possible pairwise sample comparison included in the f3-statistics test (N=8515). Subsequently, we carried out hierarchical clustering using the function *hclust* from the *R* package *stats* [62]. As input we used a distance matrix generated from the f3-statistics-derived z-scores (Fig. 1A).

Additionally, we determined the level of population structure using genetic distances coupled with dimensionality reduction methods. We calculated pairwise Hamming distances using *Plink V*.*1*.*9* [63]. Such distances were used as input for Principal Component Analysis (PCA) using the function *prcomp* from the *R* package *stats* [62] (Supp. Fig. 1A). To assess the robustness of the clusters, PCA coordinates were used to compute silhouette scores using the function *silhouette* from the *R* package *cluster [64]*. We calculated mean silhouette scores for different number of clusters (K=2-6) and found that the highest mean silhouette scores were obtained when K=4. We also used Discriminant Analysis of Principal Components (DAPC) [65], implemented in the *adegenet R* package. The analysis was carried out by capturing the variance in the 10 first PC’s. The Bayesian Information Criterion (BIC) indicated that the best number of groups was K=4 (Supp. Fig 1C-D). We used the grouping of individuals in four clusters for subsequent analyses.

### Population genetics analyses

We constructed a neighbor network using the program *SplitsTree V*.*4*.*14*.*6 [35]*. As a proxy for recombination within each of the clusters, we used the four-gamete test [38] as implemented in *RminCutter* [66]. To this aim, we created consensus *fasta* sequences from the contigs 1 to 7 using the filtered vcf file with *bcftools* V. 1.3.1 [67]. The summary statistic was calculated by dividing the total number of violation events of the four-gamete test by the total number of SNPs. Nucleotide diversity (*π*), Fixation Indices F_ST_ and Tajima’s *D* values were calculated using *vcftools V*.*0*.*5*.*15* [68].

We computed *D*-statistic values [43] as:

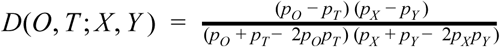

where P_O_, P_T_, P_X_, and P_Y_ are frequencies of randomly selected alleles in populations (O)utgroup, (T)est, X, and Y at each locus. The reported 95% confidence intervals were calculated as *D*±(*SE* ×1.96) where the Standard Error was computed using a jackknife weighted by the number of SNPs for each 5 Mb block in the genome [69]. We performed the calculations using *popstats* [70].

### Genomic segmentation analysis

Based on the *D*-statistic results, two isolates from the diverse group I (CH1016 and HB-LTH18) showed genome-wide introgression evidence with the clonal lineage II. In order to identify which regions of the genomes of CH1016 and HB-LTH18 show higher nucleotide similarity to clonal lineage II than to members of the diverse group I, we performed a window-based similarity analysis. These regions, especially if they overlap between CH1016 and HB-LTH18, will be strong candidates for being introgressed from the clonal lineage II. Consequently, we performed window-based pairwise nucleotide similarity comparisons between an example isolate of clonal lineage II (TW-PT3-1) and the two Chinese individuals (CH1016 and HB-LTH18). To this end we divided the seven chromosomes in 400 windows, each of which had the same number of SNPs. To ascertain the basal level of similarity among clonal lineage II individuals we compared our example clonal lineage II isolate TW-PT3-1 with another clonal lineage II isolate (BR0026). Finally, to ascertain the nucleotide similarity between clonal lineage II and non-introgressed individuals from the diverse group I, we compared our example clonal lineage II isolate TW-PT3-1 with diverse group I isolates CH0532 and CH0333.

### Bayesian tip-dated phylogenetic analysis

To perform the phylogenetic analysis, we first removed individuals from the diverse group I, as these recombining group of individuals do not comply with the assumptions of any phylogenetic analysis (Fig. 2C). We kept only biallelic variant positions to perform a Markov-Chain-Monte-Carlo-based phylogenetic reconstruction using *BEAST V*.*2*.*4*.*8* [71]. We used the isolates collection dates (Supp. Table 1) as prior information for the estimation of divergence times. We used *ModelTest-NG* [72] to assess the best suitable substitution model. Based on the lowest Akaike Information Criterion (AIC), we selected the General Time Reversible model. Since the calculation was performed with non-recombining individuals from the same species, we used a strict clock rate with a prior value of 1.98 e-8 substitutions/site/year, which was the rate ascertained in Gladieux *et al*. [30]. To test the hypothesis of a non-clocklike data, we estimated the coefficient of variation in a model relaxed clock log normal model to be 0.0042, suggesting a strong evidence for a clock-like data [46]. In order to reduce the effect of demographic history assumptions, and to calculate the dynamics of the population size through time, we also chose a Coalescent Extended Bayesian Skyline approach [73]. We combined the output of four independent MCMC chains. Each chain had a length of 10 million iterations and was logged every 1000 iterations. We only used chains with overall ESS values above 200 and summarized a maximum clade credibility tree with *TreeAnnotator*. Finally, to illustrate the impact of recombining individuals on the robustness of the phylogeny and the estimation of divergence times, we repeated the analysis including the individuals from the diverse group I.

Furthermore, to assess the divergence time between all rice-infecting clonal lineages and their closest relative, we also incorporated the *Setaria*-infecting isolate SA05-144 [24] in our analysis. We combined the output of four independent MCMC chains with the above described parameters and methodology.

### Effector genes repertoire

To determine the effector gene repertoire for each of 131 *M. oryzae* isolates described in Supp. Table 1, we mapped the publicly available genomic short read sequences from these isolates to a reference set of diverse effector coding sequences created by combining the validated effectors described in Yoshida et al. [24]and the predicted effectors reported in [50]. We reduced the redundancy of the reference by removing highly similar sequences (≥ 90% identity). The final reference set included 178 coding DNA sequences for candidate effectors (Supp. Table 5) from different *M. oryzae* lineages infecting hosts such as rice, wheat, oat, millet, and wild grasses. The coordinates of the reference effector genes corresponding to *M. oryzae* PacBio genome GUY-11 (GenBank accession GCA_002368485.1) are shown in Supplementary Table 4. We used elongation factor 2 mRNA sequence (GenBank accession XM_003714691.1) from *M. oryzae* as a positive control for presence of a gene, and a secreted protein gene *CoMC69* from the fungus *Colletotrichum orbiculare* as a negative control for absence of a gene in the reference for short read mapping.

Mapping was performed with *bwa-mem* V.0.7.15 [74]. An effector was deemed present if more than 80% of its sequence was recovered with a minimum depth of 3x, using *SAMtools* V1.6 [75].

To summarize effector content per isolate, we built a presence/absence matrix indicating presence and absence of effector genes with 1 and 0, respectively (Fig 7A-B and Supp. Table 6). For subsequent analyses we excluded effector genes that were either present or absent in all lineages, as they are uninformative for clustering algorithms. This filtering resulted in a presence/absence matrix that contains a set of 69 informative effectors. We organized the columns of this matrix according to the dendrogram of genetic groups (Fig. 1), while the rows were sorted using hierarchical clustering with the function *hclust* from *R stats* package [62].

To determine which effectors have the strongest association with the defined genetic structure of *M. oryzae*, we conducted PCA and loading analysis using the presence/absence matrix per isolate as input. Analyses were carried out with the *princomp* function of the *R stats* package [62]. Then, by multiplying the absolute value of X and Y coordinates of each loading vector in PC1 and PC2, we assessed the strength of importance per effector (Supp. Fig. 6A). We selected a subset of effectors that contains the 13 most important effectors (i.e. loading vectors with the highest magnitudes), as they together explained 90% of the variance (Supp. Fig. 6C). We recalculated the PCA using only this subset of 13 effectors, which resulted in an increase from 44.8% to 73.8% (total increase of 29%) of the variance explained by PC1 and PC2 together.

A similar analysis was also carried out using the Extremely Randomized Trees algorithm implemented in the *Python scikit-learn* module [76], with 100 trees per forest, and trained using all the effector presence/absence data. The feature importances were extracted from the trained model. This process was repeated 2,500 times to ensure consistency, and the mean effector importance for reconstructing the population structure was calculated in order to rank the effectors. Using this method, 90% of the variance was explained by 16 effectors (Supp. Fig. 6C).

### Code and data availability

Scripts and supporting data can be found at: https://gitlab.com/smlatorreo/genetic_history_of_rice-infecting_magnaporthe_oryzae

## Acknowledgements

This work was supported by the Max Planck Society and its Presidential Innovation Fund, the Gatsby Charitable Foundation, the ERC (proposal 743165), and BBSRC. We thank Michael Dannemann, and members of the Kamoun and Burbano laboratories for useful discussions and input on data analysis; Talia Karasov and Francois Balloux for comments on the manuscript, and Detlef Weigel for supporting Sergio Latorre’s visit to London during the final stages of the project.

## Authors’ contributions

SML, SK and HAB conceived the study. SML, CSR, AM, JW, SK, HAB designed and performed data analyses. SML, SK and HAB wrote the paper. All authors read and approved the final manuscript.

## Supplementary figures

**Supplementary Figure 1.**
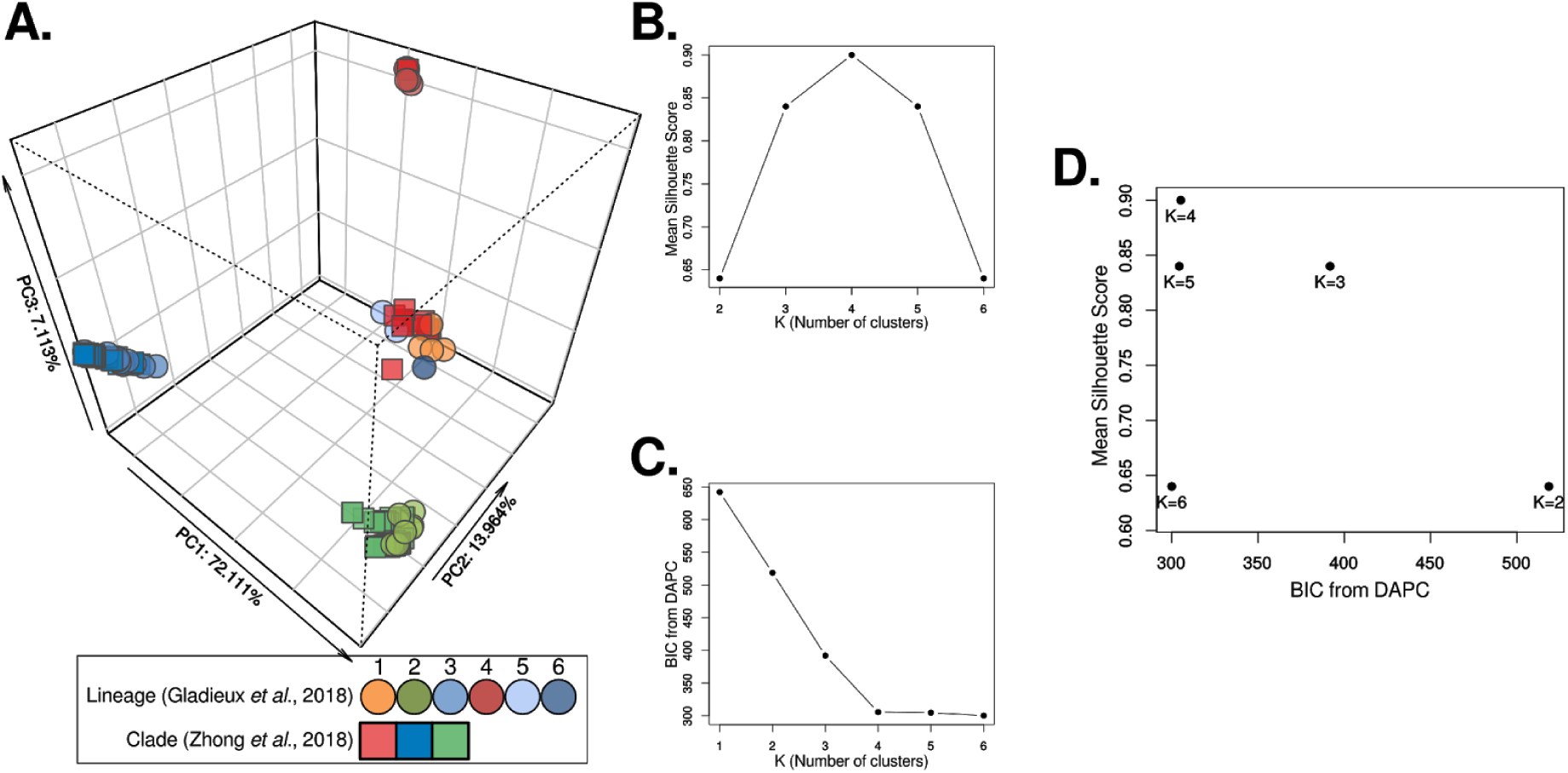
Principal component analysis (PCA) reveals four defined groups. **(A)** PCA based on pairwise Hamming distances. **(B)** Silhouette score analysis shows best averages per group scores when *K=*4. (**C**) Discriminant Analysis of Principal Components (DAPC) shows stabilization of the Bayesian Information Criterion (BIC) when *K*=4. **(D)** BIC from DAPC versus mean silhouette scores shows an optimal number of groups when *K*=4.

**Supplementary Figure 2.**
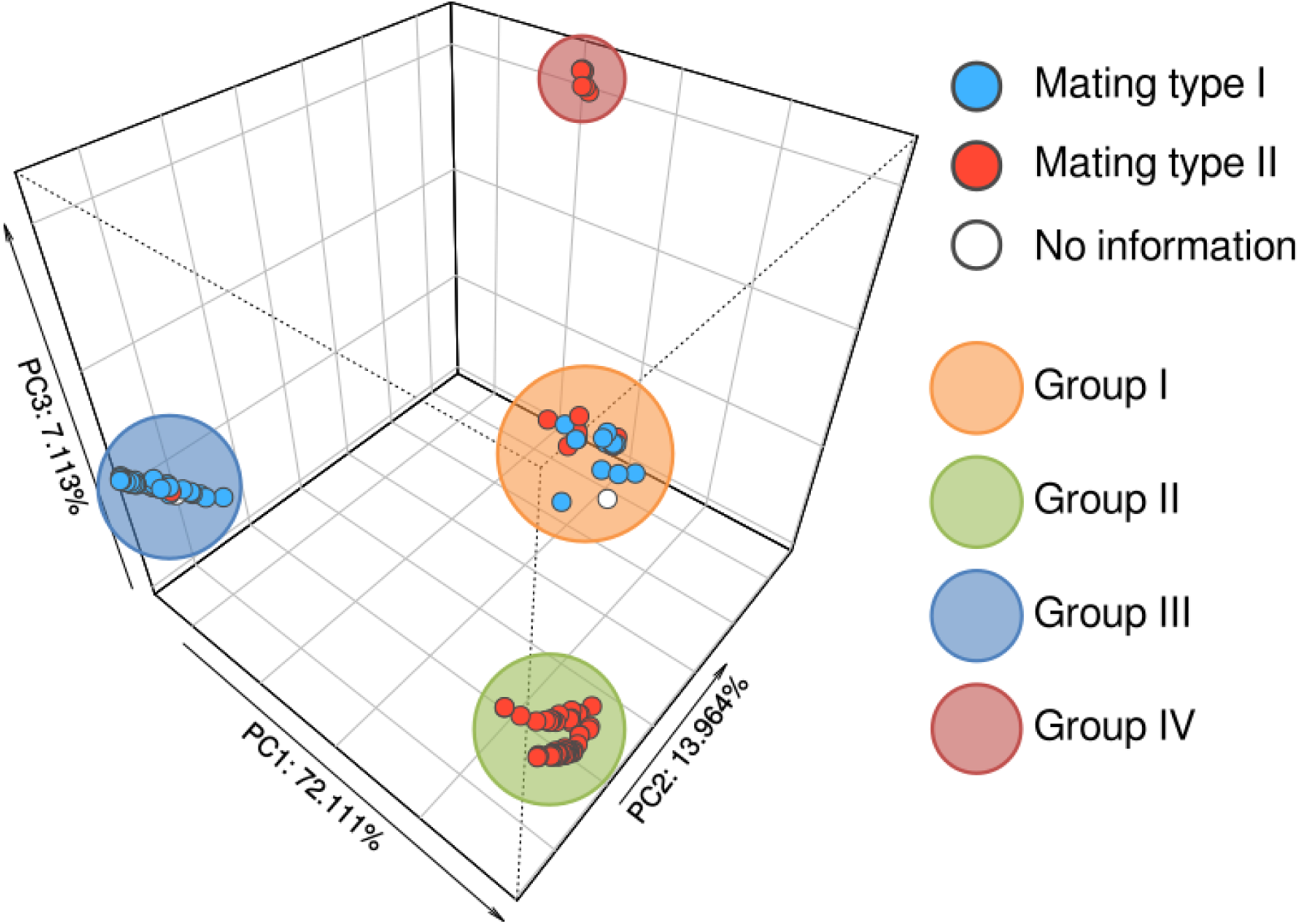
Relation between genetic groups and sample mating type. PCA based on pairwise Hamming distances (same coordinates as Supp. Fig. 1). The color scheme codifies the mating type described as metainformation in the previous studies.

**Supplementary Figure 3.**
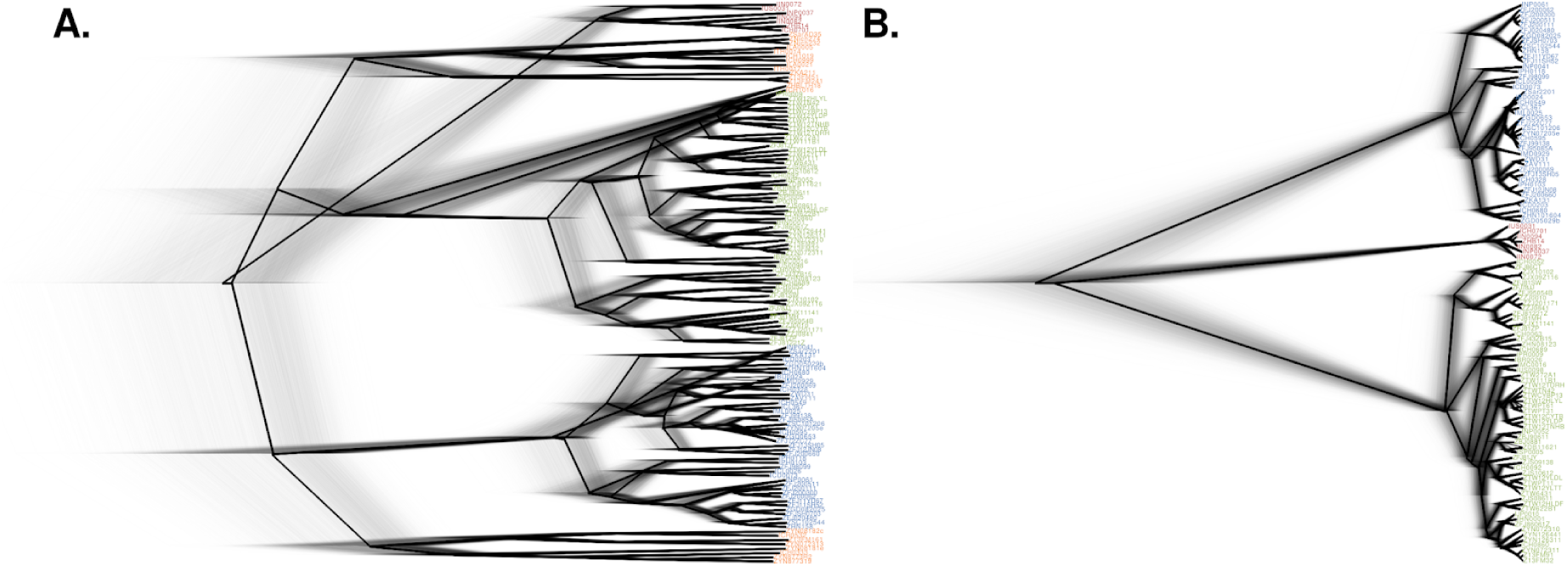
Effect of recombination on the *Magnaporthe oryzae* phylogeny construction. Comparison between phylogenies with **(A)** and without **(B)** the diverse recombining group. Grey diffused lines depict all calculated trees, whereas the black lines represent the maximum clade credibility tree.

**Supplementary Figure 4.**
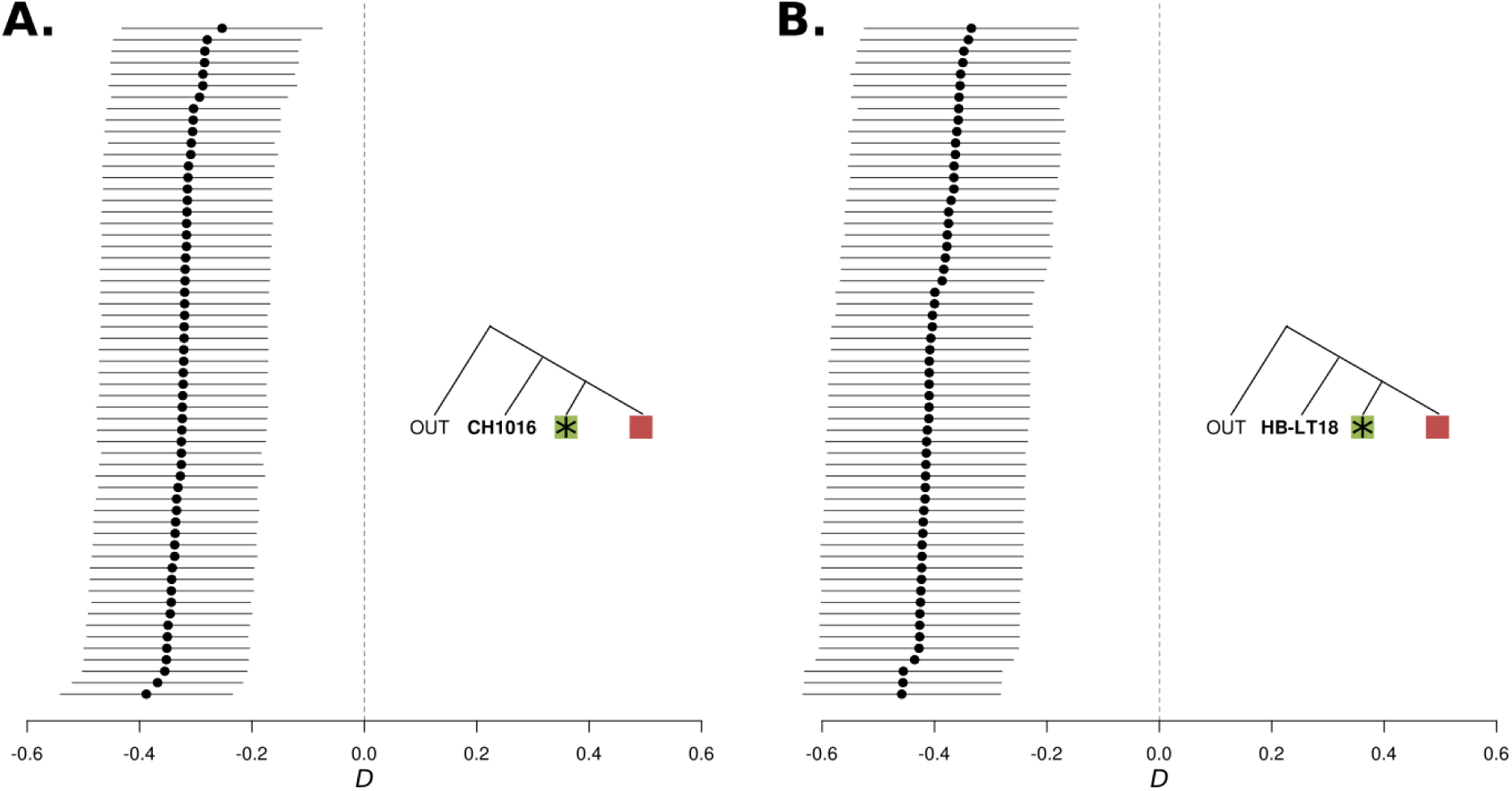
Two Chinese individuals display consistent introgression with the clonal lineage II. *D*-statistics using different phylogenetic configurations depicted as colored inset trees. The green box with the asterisk represents a position in which all individuals from the clonal lineage II were placed in an iterative way. Red boxes represent a fixed individual from the clonal lineage IV. **(A)** *D*(Outgroup, CH1016, Lineage II individual, Lineage IV individual). **(B)** *D*(Outgroup, HB-LT18, Lineage II individual, Lineage IV individual). Points represent *D*-statistic tests, and lines depict 95% confidence intervals.

**Supplementary Figure 5.**
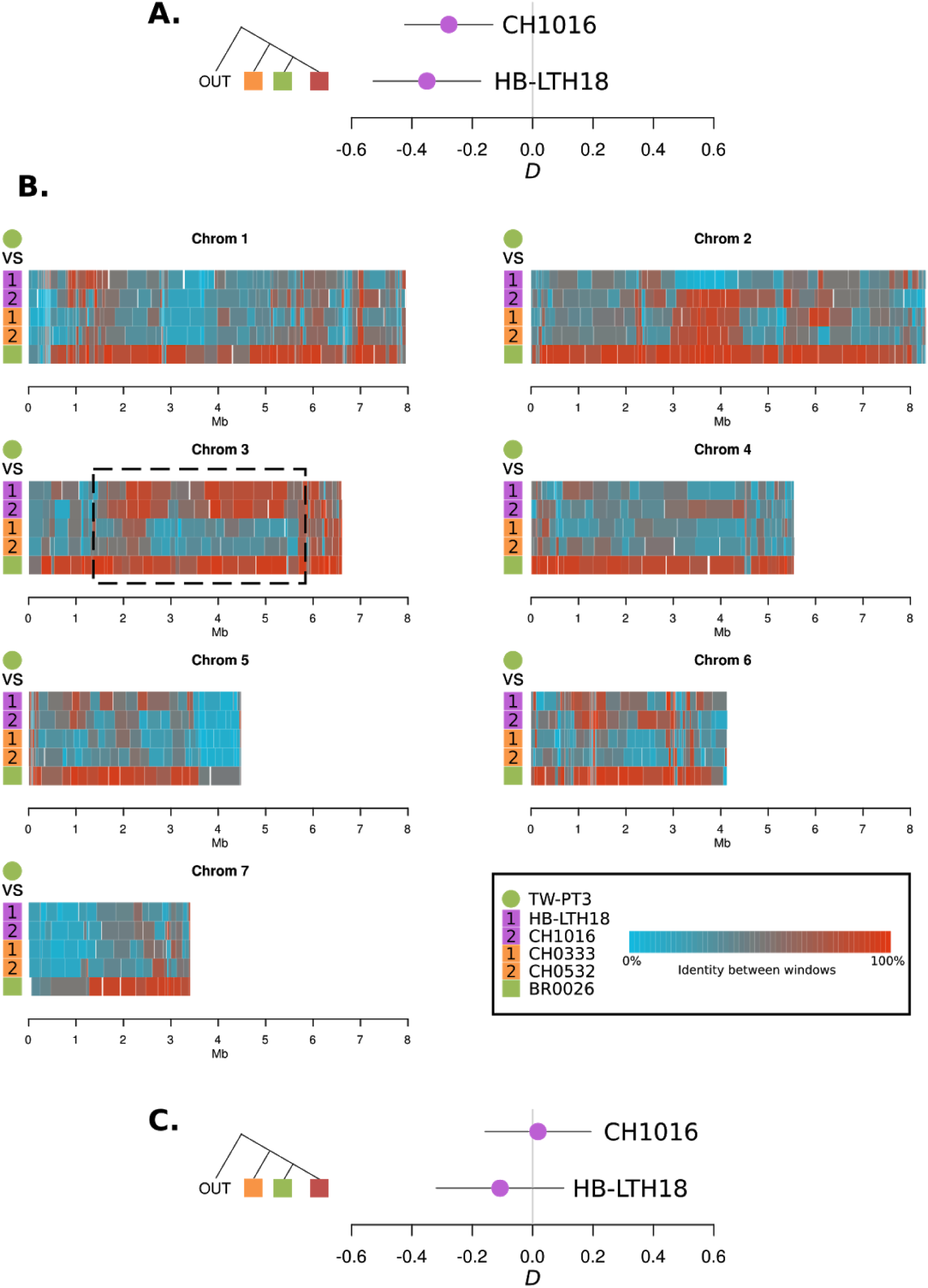
Ancestry-based genomic segmentation of Chinese individuals *CH1016* and *HB-LTH18* reveals a 4 Mb putative introgressed region on chromosome 3. **(A)** The inset tree shows the *D*-statistics configuration, *D*(Outgroup, Orange; Green, Red), used to detect introgression between clonal lineage II and two individuals (*CH1016* and *HB-LTH18*) from the diverse group I (as in Fig 5B). Introgression is inferred based on the significant negative *D*-statistics. **(B)** Each panel shows homologous chromosomes from *CH1016* and *HB-LTH18* in addition to the control individuals from the diverse group I (CH033 and CH0532) and the clonal lineage II (BR0026), segmented according to their ancestry. The color coding represents the level of SNP similarity between each individual and the chosen clonal lineage II individual (TW-PT3) for that particular segment. Chromosome 3 shows a 4 Mb segment inferred to be introgressed between the clonal lineage II and both *CH1016* and *HB-LTH18* (boxed area). **(C)** The same *D*-statistic test as in A. was carried out after all putative introgressed fragments (in red) with a percentage similarity value of >= 60 were removed. The test was not significant, i.e., no different from zero.

**Supplementary Figure 6.**
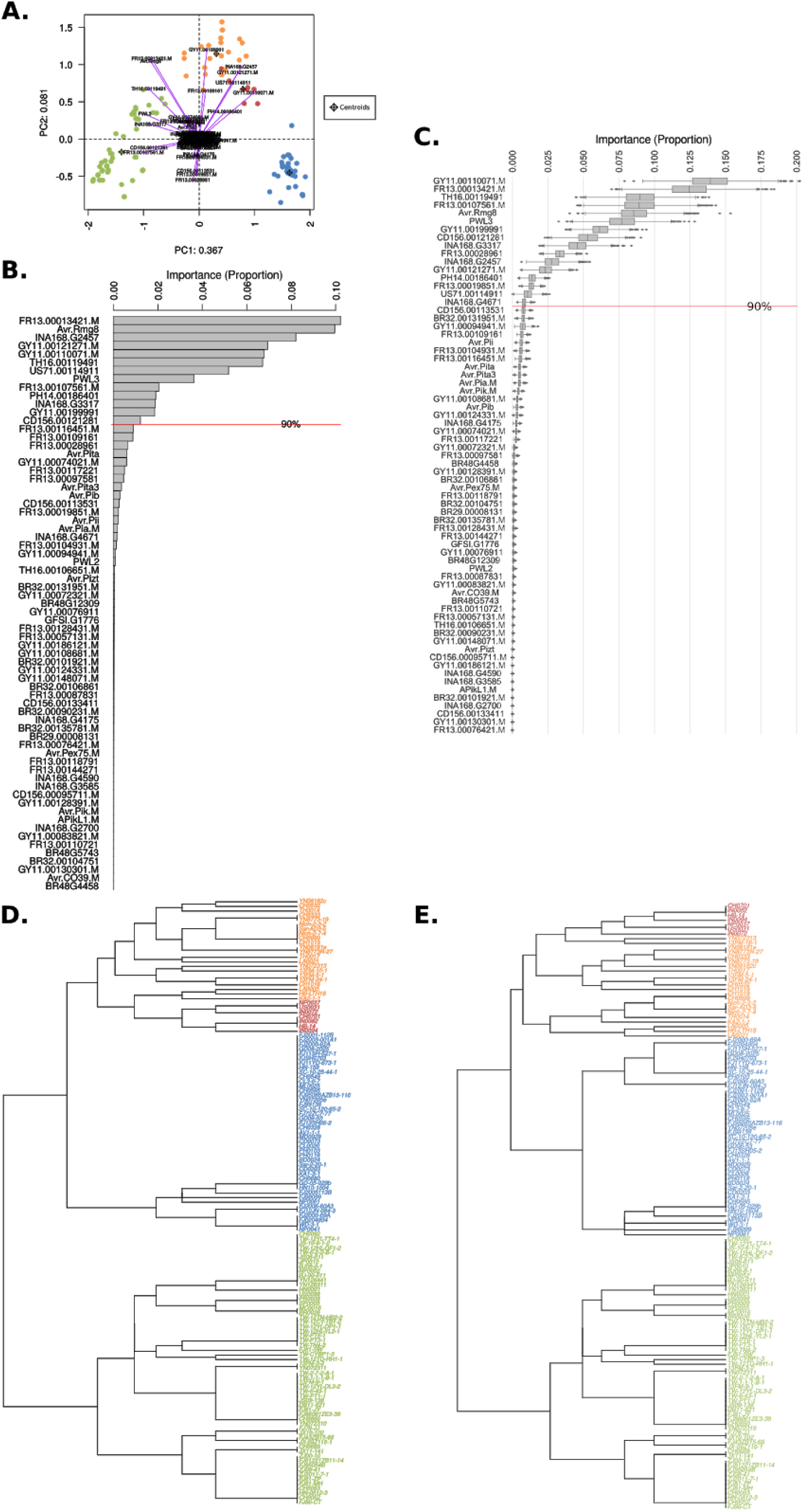
Effector loadings reveal major effector lose in clonal lineage III. **(A)** Biplot based on the presence/absence effector matrix (Fig 7C). Dots represent isolates color-coded by their genetic group. Vectors and labels correspond to the effector loadings for PC1 and PC2. All vectors were scaled by 3X for better representation. **(B)** The barplot represents the absolute value of the product of the PC1 and PC2 coordinates for each loading vector. The horizontal red line represents the cumulative sum of 90% of the data. **(C)** The set of boxplots summarize the distribution of the importance of each effector as decision factors for the genetic group assignment under 2,500 iterations of the extremely randomized trees algorithm. The horizontal red line represents the cumulative 90% of the data based on the mean values. **(D)** Hierarchical cluster-based dendrogram built by subsetting the 13 first effectors showed in B. **(E)** Hierarchical cluster-based dendrogram built by subsetting the 16 first effectors showed in **C**.

