## Supplementary material for "Recently expanded clonal lineages of the rice blast fungus display distinct patterns of presence/absence of effector genes": Supp. Table 3

**Supplementary Table 3**. Members of the pan-effectorome that are present or absent in all 131 *Magnaporthe oryzae* isolates used in this study along with the effectors showing presence/absence polymorphism

| **Present in all isolates**  **(65 effectors)** | | **Absent in all isolates**  **(44 effectors)** | **Effectors showing presence/absence polymorphism (69 effectors)** | |
| --- | --- | --- | --- | --- |
| Avr.Pi9 | FR13.00107801.M | APikL3.M | APikL1.M | FR13.00116451.M |
| BR48G12130 | FR13.00108711.M | DIG41.G5801 | Avr.CO39.M | FR13.00117221 |
| BR48G630 | FR13.00114581.M | DIG41.G7456 | Avr.Pex75.M | FR13.00118791 |
| BR48G9741 | FR13.00122001.M | BR29.00004921.M | Avr.Pia.M | FR13.00128431.M |
| BR58.G2338 | FR13.00143751.M | BR29.00026641.M | Avr.Pib | FR13.00144271 |
| BR58.G4698 | GY11.00054591.M | BR29.00027131.M | Avr.Pii | GY11.00072321.M |
| TH16.00100711.1.M | GY11.00060011 | BR29.00043011.M | Avr.Pik.M | GY11.00074021.M |
| BR32.00003601.M | GY11.00067931 | BR29.00052161.M | Avr.Pita | GY11.00076911 |
| BR32.00040381 | GY11.00069551.M | BR29.00081821.M | Avr.Pita3 | GY11.00083821.M |
| BR32.00066181.M | GY11.00096851 | BR29.00089541 | Avr.Pizt | GY11.00094941.M |
| BR32.00081411.M | GY11.00128491.M | BR29.00091361.M | Avr.Rmg8 | GY11.00108681.M |
| BR32.00101991.M | GY11.00137731.M | BR29.00091381.M | BR48G12309 | GY11.00110071.M |
| BR32.00117431.M | GY11.00138681.M | BR29.00091681.M | BR48G4458 | GY11.00121271.M |
| BR32.00131211.M | TH12.00085091.M | BR29.00092011 | BR48G5743 | GY11.00124331.M |
| BR32.00132551 | TH12.00121641.M | BR29.00094821.M | GFSI.G1776 | GY11.00128391.M |
| BR32.00133701.M | TH16.00079081.M | BR29.00096171.M | INA168.G2457 | GY11.00130301.M |
| BR32.00141021.M | TH16.00118381.M | BR29.00096401 | INA168.G2700 | GY11.00148071.M |
| BR32.00141031.M | US71.00065681.M | BR29.00096771.M | INA168.G3317 | GY11.00186121.M |
| BR32.00141181.M | US71.00116921.M | BR29.00097421 | INA168.G3585 | GY11.00199991 |
| CD156.00118721 | MGDIG41 | BR29.00105021.M | INA168.G4175 | PH14.00186401 |
| CD156.00122701.M | MOBR58 | BR29.00105921 | INA168.G4590 | TH16.00106651.M |
| CD156.00131711.M |  | BR29.00106461.M | INA168.G4671 | TH16.00119491 |
| CD156.00138051.M |  | BR29.00107161 | BR29.00008131 | US71.00114911 |
| FR13.00001281.M |  | BR29.00107301 | BR32.00090231.M | PWL2 |
| FR13.00003821.M |  | BR29.00112111.M | BR32.00101921.M | PWL3 |
| FR13.00004061.M |  | BR29.00114891.M | BR32.00104751 |  |
| FR13.00020841.M |  | BR29.00115871 | BR32.00106861 |  |
| FR13.00035971 |  | BR29.00115881 | BR32.00131951.M |  |
| FR13.00041321.M |  | BR29.00117591.M | BR32.00135781.M |  |
| FR13.00057121.M |  | BR29.00118801.M | CD156.00095711.M |  |
| FR13.00066391.M |  | BR29.00119471 | CD156.00113531 |  |
| FR13.00067231 |  | BR29.00119491.M | CD156.00121281 |  |
| FR13.00068741.M |  | BR29.00119511.M | CD156.00133411 |  |
| FR13.00069491 |  | BR29.00121041.M | FR13.00013421.M |  |
| FR13.00070821.M |  | BR29.00121721 | FR13.00019851.M |  |
| FR13.00080861.M |  | BR29.00125251.M | FR13.00028961 |  |
| FR13.00081261.M |  | BR32.00090371.M | FR13.00057131.M |  |
| FR13.00088361.M |  | BR32.00101931.M | FR13.00076421.M |  |
| FR13.00089161.M |  | BR32.00101951.M | FR13.00087831 |  |
| FR13.00093031.M |  | BR32.00102581 | FR13.00097581 |  |
| FR13.00094721.M |  | BR32.00120141 | FR13.00104931.M |  |
| FR13.00094761 |  | BR32.00128081.M | FR13.00107561.M |  |
| FR13.00099751.M |  | BR32.00143481 | FR13.00109161 |  |
| FR13.00101061 |  | US71.00000121.M | FR13.00110721 |  |
