## Supplementary material for "Recently expanded clonal lineages of the rice blast fungus display distinct patterns of presence/absence of effector genes": Supp. Table 4

**Supplementary Table 4**. Patterns of effector presence/absence polymorphism in *Magnaporthe oryzae* isolates

| **Pattern** | **Group showing presence** | **Group showing absence** | **Effector** |
| --- | --- | --- | --- |
| Lineage/group specific | I, II, IV | III | TH16.00119491*  FR13.00013421.M*  Avr-Rmg8*  CD156.00121281* |
|  | I, II | III, IV | PWL3*  INA168.G3317*  FR13-00107561.M* |
|  | I, II, III | IV | FR13.00028961* |
|  | I, III, IV | II | GY11.00110071.M* |
|  | I | II, III, IV (with a few exceptions) | GY11.00199991* |
| Patchy distribution | Varies | Varies | Avr-Pia.M  Avr-Pii |
| Partial gene loss within a group | I, III, IV | II | PH14.00186401*  GY11.00094941.M  GY11.00121271.M*  INA168.G2457*  US71.00114911* |
| Near-complete gene loss across all groups | Varies | Varies | FR13.00109161  APikL1  CD156.00133411  Avr-CO39.M  BR32.00090231.M  BR29.00008131  GFSI.G1776 |

*These are part of top 13 (as well as top 16) effectors used in Supp. Fig. 5B-E.
